# Evolution and integration of a novel cell type in the housefly Love Spot

**DOI:** 10.64898/2026.06.23.734072

**Authors:** Antoine Donati, Yunchong Zhao, Eleanor Terner, Nikos Konstantinides, Michael W. Perry

## Abstract

The addition of new neuron types is thought to underlie the evolution of complex nervous systems, yet the mechanisms by which they arise remain unclear. Here we investigate the evolution of a novel target detection photoreceptor in the housefly *Musca domestica*. In males, a dorso-frontal eye region known as the “Love Spot” enables rapid detection and tracking of females during mating flights. In this region, R7 photoreceptors - normally dedicated to color vision - are repurposed for target detection through altered Rhodopsin expression, physiology, and circuit connectivity. We show that these Love Spot R7 cells (LsR7) are initially specified as canonical R7 photoreceptors but later adopt a chimeric identity combining transcription factors normally associated with either color or motion vision. The sex-determination transcription factor Doublesex (Dsx) is strongly upregulated after R7 specification and, together with the transcription factor Spineless, drives the LsR7 gene regulatory program. Because Dsx upregulation occurs after initial axon targeting to the medulla, LsR7 axons subsequently shorten to connect with lamina neurons and OFF-pathway motion vision circuits. These findings show how new combinations of cell-identity regulators can generate novel neural cell types while revealing developmental constraints imposed by evolutionary history.

## Main

The extraordinary diversity of neuron types in animal brains was first highlighted by Santiago Ramón y Cajal in the late 1800s^1^ after a new single-neuron staining technique enabled beautiful drawings of their diverse morphology^2^. This stunning variety has been challenging to fully characterize in any single species, yet also evolves across species. In many lineages, nervous systems have gained complexity through the expansion of total neuron numbers and the diversification of specialized cell types^3^. Over time, novel neurons have evolved to enable specialized behaviors^2–5^. How specific genetic changes can produce new types of neurons is largely unknown.

New tools have made it possible to profile gene expression in large numbers of cell types in entire tissues^6^ and to begin to match those expression patterns with neurons of specific morphologies. Cell atlas projects have characterized neural diversity in species ranging from *Drosophila melanogaster*^7^, to mice^8^, to humans^9^. In parallel, other studies have sought to identify the developmental and genetic basis of how such an extraordinary diversity of cell types is produced^10–17^. These efforts have uncovered general mechanisms responsible for generating neural diversity, including the tightly choreographed integration of spatial and temporal cues. The outcome of this process is the expression of cell-type specific combinations of transcription factors. These networks of interacting transcription factors then orchestrate the generation of neuron-specific features such as precise morphology, connectivity, and physiology^15,18^. How this combinatorial system that specifies diverse cell fates has been modified to give rise to even more cell types, and how new cell types integrate into existing neural circuits to provide novel function, is largely unknown.

Comparisons between evolutionarily distant model species have provided limited insight into how new neurons first evolve, yet identifying novel neural types between closely related species, such as between *Drosophila* species, can also be challenging. To address this, we examined an intermediate evolutionary distance and focused on a specialized cell type within the ’Love Spot’ region of the male *Musca domestica* (housefly) retina. Sexual selection is a major source of morphological and behavioral diversity in animals, and sexually dimorphic traits are gained and lost rapidly during evolution^19,20^. The Love Spot is one of the most striking sexually dimorphic traits in insects (**Fig1a**). This modified dorsal region of the male eye is used to detect and chase females in flight and has a long history of study in houseflies^21,22^. Similar features have been observed in many dipteran families^23^, though Love Spot-like modifications have a sparse phylogenetic distribution, are correlated with aerial pursuit, and are absent in *D. melanogaster*^21,22^. The most obvious feature is the enlargement of dorsal ommatidia (unit eyes) in males, often combined with the two eyes sitting closer together dorsally (**Fig1b**). Enlarged facets in this region increase photon capture, while reduced interommatidial and acceptance angles together produce a finer-grained, higher resolution image of the dorsal visual field, preserving the contrast of a small female against the bright background of the sky. In houseflies, Love Spot photoreceptor physiology is tuned to detecting flying females^24,25^. Males perform frequent long duration chases and keep their target in their dorsal field of view while pursuing it from below^26^ (**Fig1c**), consistent with a dorsal specialization for aerial pursuit. Within this heavily modified region, R7 photoreceptors that normally serve color vision are instead repurposed for target detection and tracking^27–29^.

### The LsR7 as a novel cell type

Each ommatidium (unit eye) of a fly eye is composed of 20 cells including cone cells, pigment cells, and eight photoreceptors (R1-8), which are sensitive to different wavelengths of light depending on which Rhodopsin they express (**Fig1d**)^21^. In *D. melanogaster*, “outer” photoreceptors R1-6 express broad-spectrum Rhodopsin 1 (Rh1) and are used in motion vision while the central “inner” photoreceptors R7 and R8 sit on top of one another and make color comparisons^30,31^. As in *D. melanogaster*, the housefly eye is composed of a stochastic mosaic of two main ommatidial subtypes called “yellow” and “pale”, which are present in a 70/30 ratio^28,32^ (**Fig1e**). These were named for having either yellow or no color (pale) in their central R7 when viewed with white transmitted light^33^.

Ommatidia within the *M. domestica* dorso-frontal Love Spot region contain a novel type of R7 photoreceptor with unique characteristics, the Love Spot R7 (LsR7)^34^ (**Fig1e**). Although these LsR7 have the same position within the ommatidia as other R7s, when viewed under blue light illumination, they produce the same reddish fluorescence as outer photoreceptors, suggesting that they might instead play a role in motion vision^34^. It was also shown through intracellular recordings that LsR7 have similar broad-spectrum sensitivity as R1-6^28^ and that their axons project to the lamina, like motion-vision outer photoreceptors, instead of the medulla like all other R7 color photoreceptors^29,34^(**Fig1f_i_**). In the lamina, LsR7s axons stop in the superficial layer and form a characteristic 10µm-long right-angled branch (**Fig1f_ii_**) that makes gap junctions connections with outer photoreceptors and R8^28,29,34^. This unusual axon terminal synapses specifically on lamina neurons L2 and L3 but not L1, unlike R1-6^29^. Studies in *Drosophila* have shown that L1 neurons respond to light increments (ON pathway) while L2 and L3 respond to light decrements (OFF pathway)^35–38^. This suggests LsR7 neurons specifically send input to the OFF pathway, which we propose fits their function in detecting small dark objects (like females) against the bright background of the sky. We set out to further characterize phenotypic differences that set LsR7s apart from other R7s and to investigate the gene regulation underlying their differentiation during eye development.

### Characterizing a novel neuron

#### LsR7 identity

We first characterized Rhodopsin expression in *M. domestica*. In *D. melanogaster*, yellow R7 (yR7) express Rh4 and pale R7 (pR7) express Rh3. However, an Rh4 ortholog is missing from *M. domestica* (Rh4 was lost in the common ancestor of Muscidae, Calliphoridae and Glossinidae)^39,40^. Immunostaining with antibodies we had generated (see Methods) showed that *M. domestica* instead express Rh2, a Rhodopsin expressed only in ocelli in *D. melanogaster*, in yR7s (**Fig1g**). Like *D. melanogaster*, *M. domestica* express their yR7 and pR7 Rhodopsins in a 70/30 ratio^28,32^. In female retinas and male retinas outside the Love Spot region, Rhodopsin expression was otherwise like *D. melanogaster*, with Rh6 in R8 matched with Rh2 (instead of Rh4) in R7, and Rh5 (R8) matched with Rh3 (R7)(**Extended Data Fig1a,b**).

In the Love Spot region, both Rh2 and Rh3 expression were absent. Previous measurements found that LsR7 spectral sensitivity is similar to Rh1-expressing outer photoreceptors^28^. Immunostaining indeed revealed that LsR7 express Rh1 (**Fig1h**). Similarly, immunostainings revealed that R8s in the Love Spot region express neither Rh5 nor Rh6 (**Extended Data Fig1a**). Based on electrophysiological recordings these likely express Rh1^28^. Thus, *M. domestica* female retinas are composed of a mosaic of randomly arranged yellow and pale ommatidia similar to *D. melanogaster*, whereas males have an additional homogeneous ommatidial type in the Love Spot region in which LsR7 and R8 both express Rh1 (**Fig1i**).

To further characterize LsR7 neurons, we looked at their axonal projections in the optic lobe. LsR7 axons terminate in the superficial layer of the lamina as opposed to the medulla^29,34^. Using a *D. melanogaster* antibody for the photoreceptor-specific adhesion molecule FasII, we confirmed that LsR7 axon terminals were indeed missing from the medulla in a region that corresponds to the Love Spot in dorsal male adult optic lobes, whereas LsR8 axons that terminate in a more superficial layer remained visible (**Fig1j**). These striking changes prompted us to investigate the gene regulatory network underlying the development of this male-specific cell type.

#### Molecular characterization and chimeric identity

We next used single cell RNA sequencing (scRNAseq) to characterize gene expression in developing LsR7s. Previous work in *D. melanogaster* has shown that transcriptional diversity during visual system development peaks at mid pupation (P50), facilitating the separation of cell types^13,14,41^. We thus performed scRNAseq on *M. domestica* male and female P50 dissected retinas (**Fig2a**). We identified photoreceptor clusters using the marker genes *elav* (neural), o*rthodenticle* (o*td*) (all photoreceptors, R1-8), *spalt major* (*salm*) (R7-8), and *defective proventriculus* (*dve*) (R1-6) (**Extended Data Fig2a**). Other clusters identified included glia (*repo* or *wrapper*), muscle cells (*myosin* and actin-related genes such as *zasp66*), cone cells (*cut*), and pigment cells (*lozenge*) (**Extended Data Fig2a**).

To identify photoreceptor subtypes, including LsR7, we subsetted and re-clustered photoreceptors only (**Fig2b**). Using marker genes known from work on *D. melanogaster* photoreceptor development, we identified photoreceptor subtypes (**Fig2b,d, Extended Data Fig3**): outer photoreceptors R1-6 (*dve*) separated into an R1+R6 cluster (*bar-H1*), two clusters corresponding to R3+R4 (*sevenup* (*svp)-positive, bar-H1 negative*), and R2+R5 (*dve*-positive, *svp* and *bar-H1* negative). Inner photoreceptors expressed *salm* (along with *spalt related*, *salr*, together referred to as “*sal*”) and included R8 (*senseless (sens)*), Dorsal rim area (DRA) R7/R8 (*homothorax* (*hth*)), yR7 (*prospero (pros)*, *spineless* (*ss*)), pR7 (*pros*, *runt* (*run*)) and a small LsR7 cluster, present only in males (**Fig2b** arrow). Differential expression analysis between LsR7 and other R7 types (yR7 and pR7) revealed that the transcription factor *doublesex* (*dsx*), which is one of the key genes involved in insect sexual dimorphism^42^, was highly upregulated in LsR7 (**Fig2b,d**). Interestingly, the yR7 and LsR7 clusters were linked by another cluster of *pros*- and *ss*-expressing photoreceptors that we hypothesized could be young yR7s (corresponding to the anterior retina, where cells are younger than in the posterior retina). Indeed, pseudo-time analysis revealed two differentiation trajectories from these young yR7s to either the yR7 fate or the LsR7 fate (**Fig2c**).

LsR7 expressed *pros* and *ss*, like yR7s, but strikingly, the expression level of *dve* was higher than in yR7s, similar to that seen in outer photoreceptors (R1-6), whereas *salm* and salr expression levels were intermediate between inner photoreceptors (R7-8) and outer photoreceptors (R1-6) (**Fig2d**). Sal and Dve are involved in photoreceptor differentiation in *D. melanogaster*, where Sal is required for R7-8 differentiation^43^ and Dve in Rhodopsin expression regulation in R1-6^44^. The most novel difference in transcription factor expression was the strong upregulation of *dsx*, suggesting that Dsx might regulate LsR7 fate specification directly, perhaps by repressing *sal* and upregulating *dve*. Together, these data show that LsR7s upregulate a motion vision transcription factor, Dve, while downregulating expression of a key color vision transcription factor, Sal.

### The divergence of LsR7 identity requires Dsx

To obtain a dynamic view of LsR7 patterning, we next assessed Dsx protein levels relative to other potential regulators at different time points (**Fig2e**). We made antibodies to *M. domestica* Dsx, Ss, and Run (see Methods) and these served as markers of LsR7, yR7, and pR7 fate, respectively. Because *dsx* is differentially spliced in males and females^45^, we raised a Dsx antibody against an epitope present in both the male (DsxM) and female (DsxF) Dsx proteins. In both male and female L3 eye discs (late larval stages), Dsx was expressed in a dorsal region (arrow) anterior to the morphogenetic furrow (dashed line) in undifferentiated retinal progenitor cells in the region of the future Love Spot (**Fig 2e, Extended Data Fig2b**). By P30, the morphogenetic furrow had passed through most of the Love Spot region, and Dsx remained low in outer photoreceptors in the LS region but increased in dorso-frontal R7s-slightly in females and dramatically, though variably, in males. By P50, all LsR7s expressed similar very high levels of Dsx (**Fig2e**) whereas female R7 Dsx levels remained low (**Extended Data Fig2b**). This pattern was maintained in late pupal stages (P70). Immunochemistry confirmed at the protein level what we had observed for mRNA levels in the scRNAseq data: all LsR7s expressed Ss and not Run (**Fig2f_i_**), while Sal levels decreased and Dve levels increased (**Fig2f_ii_**). This antibody stain time series, together with our scRNAseq results, suggests that LsR7s are initially patterned as normal R7s (expressing *pros* and *sal*) and that early R7s in the dorso-anterior region diverge over time to adopt the novel LsR7 identity that includes chimeric expression of the transcription factors that pattern both color (Sal, Ss) and motion detectors (Dve).

**Figure 1.**
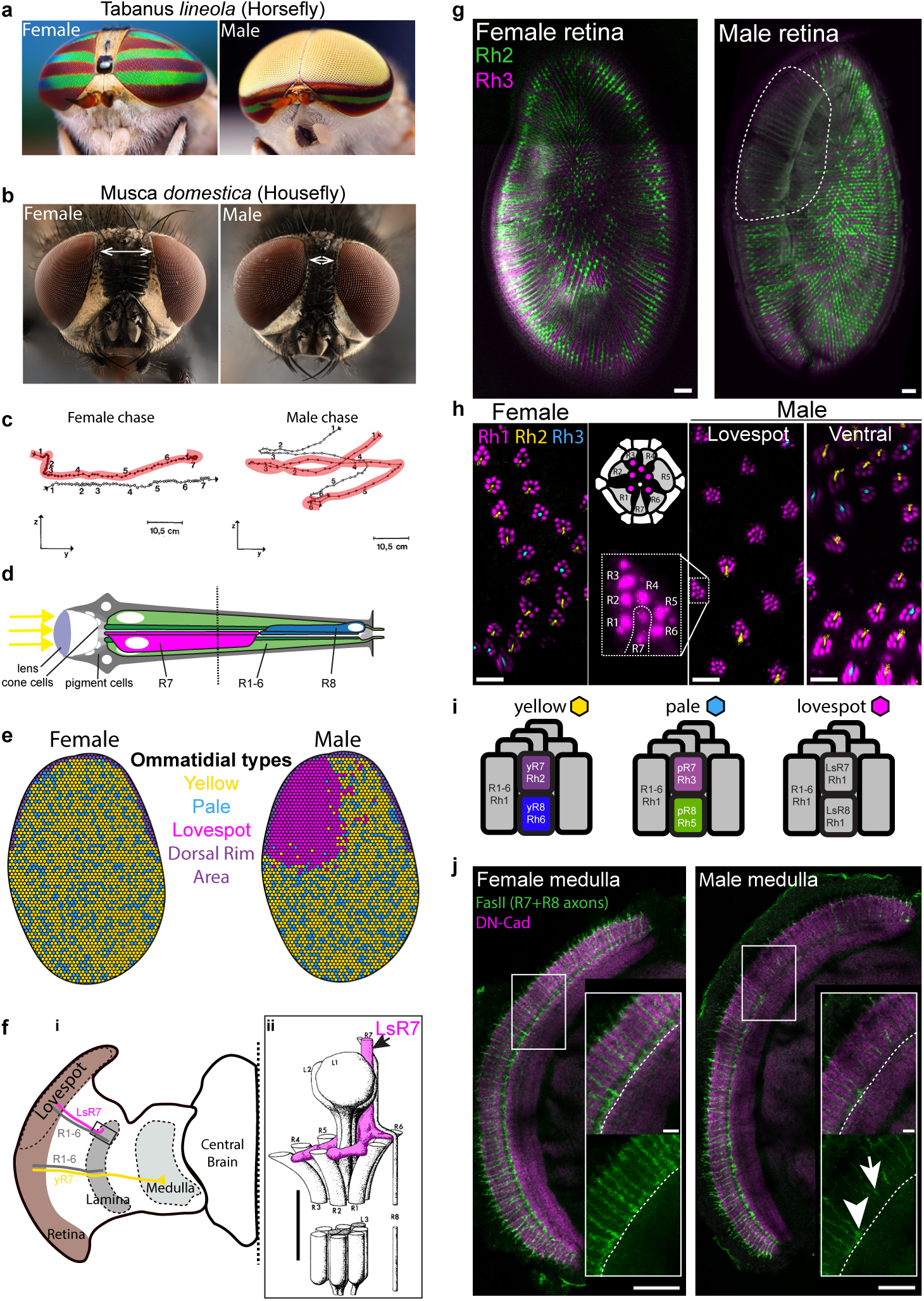
Change of Rhodopsin and axon projection of Love Spot R7 (LsR7). **a)** Many dipteran species, such as this Tabanus *lineola*, have strikingly sexually dimorphic eyes. In males, the dorsal anterior eye, or Love Spot, is highly enlarged and used to detect and pursue females in flight (image credit: Thomas Shahan). **b)** In the housefly Musca *domestica*, the Love Spot is made obvious by the smaller distance between eyes in males compared to females (double-arrows). **c)** Tracking of female or male housefly chases reveals that females chase other flies from above, using their ventral eye, whereas males chase from below using their dorsal eye. In addition, female chases are short and poorly controlled compared to males’ (adapted from^26^, chaser track is highlighted in red, time between two successive positions: 12.5ms). **d)** Each ommatidium (unit eye) is composed of a lens that focuses light on the Rhodopsin-containing rhabdoms of eight types of photoreceptors, R1 to R8. In *D. melanogaster*, inner photoreceptors R7 and R8 are mainly involved in color vision, whereas outer photoreceptors R1 to R6 are involved in motion vision. **e)** Schematics of the retinal mosaic in female and male houseflies. Female retinas are composed of a mosaic of yellow (yellow) and pale (cyan) ommatidia and polarized light-detecting dorsal rim area (DRA) ommatidia (purple) at the dorsal edge, similar to *D. melanogaster*. Male houseflies have an additional type of ommatidia in the Love Spot region (magenta). **f) (i)** Schematic of the right retina, optic lobe and central brain in male houseflies. R1-6 (grey) project their axon to the proximal lamina, yR7 (yellow) to the medulla. In contrast, LsR7 axons (magenta) stop in the superficial layer of the lamina and form a characteristic 10µm-long right-angled branch. **(ii)** Electron-microscopy-based reconstruction of the region of the superficial lamina corresponding to the box in i). The LsR7 terminal (magenta) extends perpendicular to the axon and contacts lamina neurons (L1-3) as well as outer photoreceptor (R1-6) and R8 axons (adapted from^29^). **g)** Female housefly eye R7 express Rh2 (yellow R7s, green) or Rh3 (pale R7s, magenta) in a 70/30 ratio that is similar to *D. melanogaster* Rh4/Rh3 70/30 retinal mosaic. Male housefly eye R7 display a similar Rh2/Rh3 mosaic outside the Love Spot, but do not express either of these Rhodopsin in the Love Spot (dashed-line region). **h)** Outer photoreceptors in both male and female houseflies express Rh1 (magenta), whereas R7s express Rh2 or Rh3. In the Love Spot region LsR7 express Rh1 (inset). **i)** Summary of Rhodopsin expression in different types of ommatidia in houseflies: outer photoreceptors always express Rh1, yellow ommatidia have Rh2-expressing R7 (yR7, dark purple) and Rh6-expressing R8 (yR8, blue), while pale ommatidia have Rh3-expressing R7 (pR7) and Rh5-expressing R8 (pR8, green). In Love Spot ommatidia, both Love Spot R7 (LsR7) and Love Spot R8 (LsR8) express Rh1 only. **j)** Immunostaining of photoreceptor axons with FasII (green) shows that in female houseflies, R7 axons all terminate in the same layer of the medulla, just below a dark band in the DN-Cadherin staining (magenta) that likely corresponds to the serpentine layer, located near medulla layer M6 where R7 axons terminate in *D. melanogaster*. R8 terminals are not visible because they overlap with the distal part of the R7 axons. In contrast, male housefly R7 terminals are missing in the dorsal medulla, where dorsal LsR7 should send their axons, revealing the LsR8 terminals (inset, arrow, compare with a more ventral non-Love Spot R7 terminal, arrowhead).

**Figure 2.**
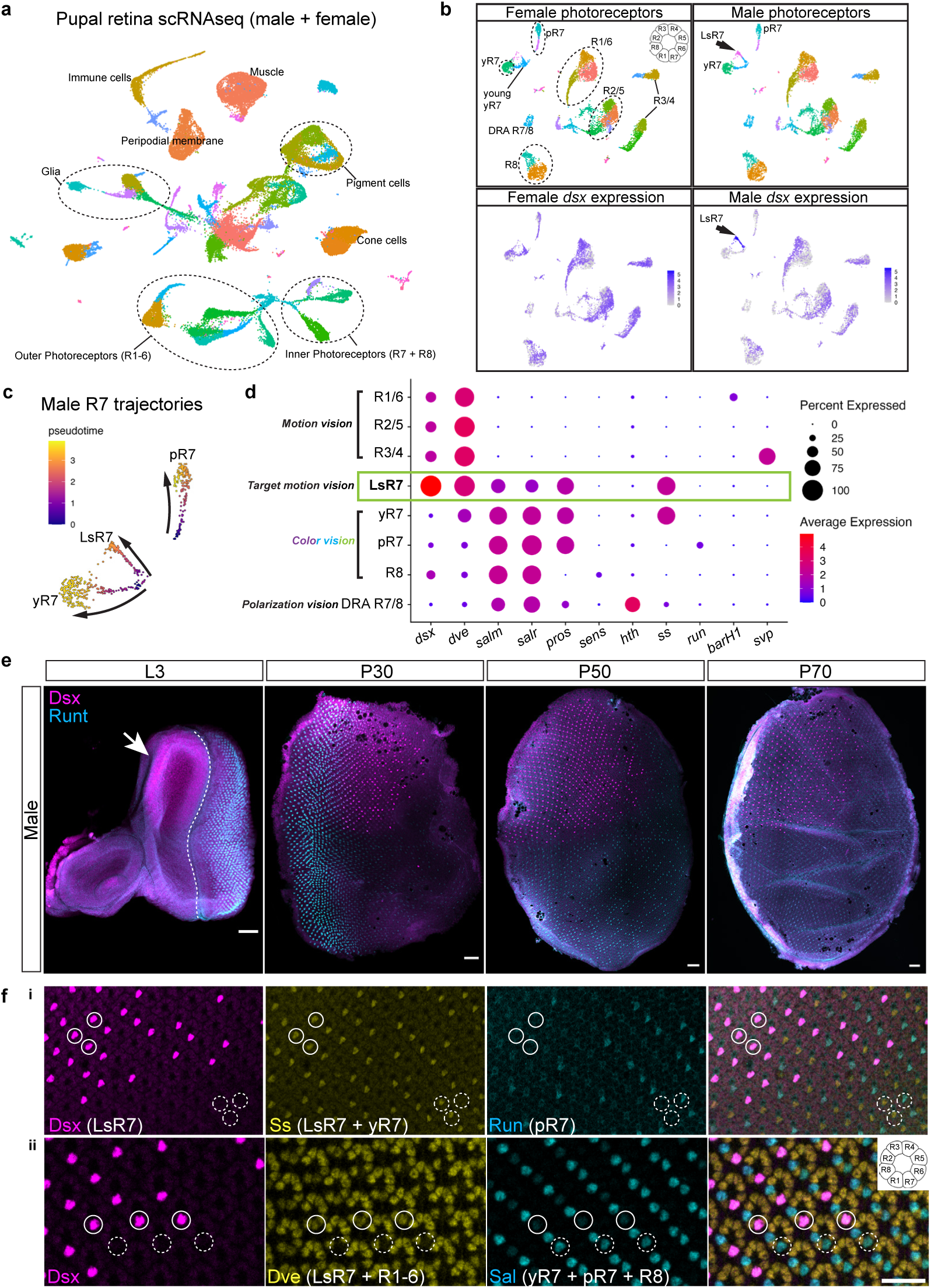
The LsR7 is a hybrid cell type expressing high levels of Dsx during development a-d) Single cell RNAseq of pupal retinas. **a)** UMAP plot of P50 male and female housefly retinas. The markers used to identify the different cell types are shown in **Extended Data Fig1**. **b)** Subsetting and re-clustering of photoreceptors reveals photoreceptor subtypes including a male-specific R7 cluster corresponding to LsR7. The inset shows the relative position of photoreceptors in a pupal retina. Pseudotime analysis reveals three differentiation trajectories for male R7: pale, yellow, and Love Spot. **c)** Dsx expression in photoreceptors. Both male and female photoreceptors express dsx. In males, LsR7 express very high levels of dsx (arrow). **d)** Expression of *dsx* and classic *D. melanogaster* photoreceptor marker genes in housefly pupal photoreceptors. LsR7 express very high levels of *dsx*, *dve* levels similar to outer photoreceptors (R1-6, motion vision) and lower levels of *salm* and *salr* compared to inner photoreceptors (R7-8, color vision). LsR7 also express *pros* (a R7 marker) and *ss* (a yR7 marker). **e)** At late larval stages (L3), Dsx is expressed in a dorsal region of the eye disk, before the morphogenetic furrow (dashed line) in males (arrow). By P30, Dsx expression is ramping up in LsR7 whereas it remains weak in other cells. At P50 and P70, Dsx levels are stable and high across all LsR7. Scale bars 50µm. **f)** Protein levels of Dsx and other photoreceptor marker genes at the edge of the Love Spot at P50. **(i)** All LsR7 (plain circles) express Ss and not Run whereas R7 outside of the Love Spot (dashed circles) express either Ss (yR7) or Run (pR7). **(ii)** All LsR7 (plain circles) have increased levels of Dve and reduced levels of Sal compared to other R7 outside the Love Spot (dashed circles). Scale bar: 25µm.

**Figure 3.**
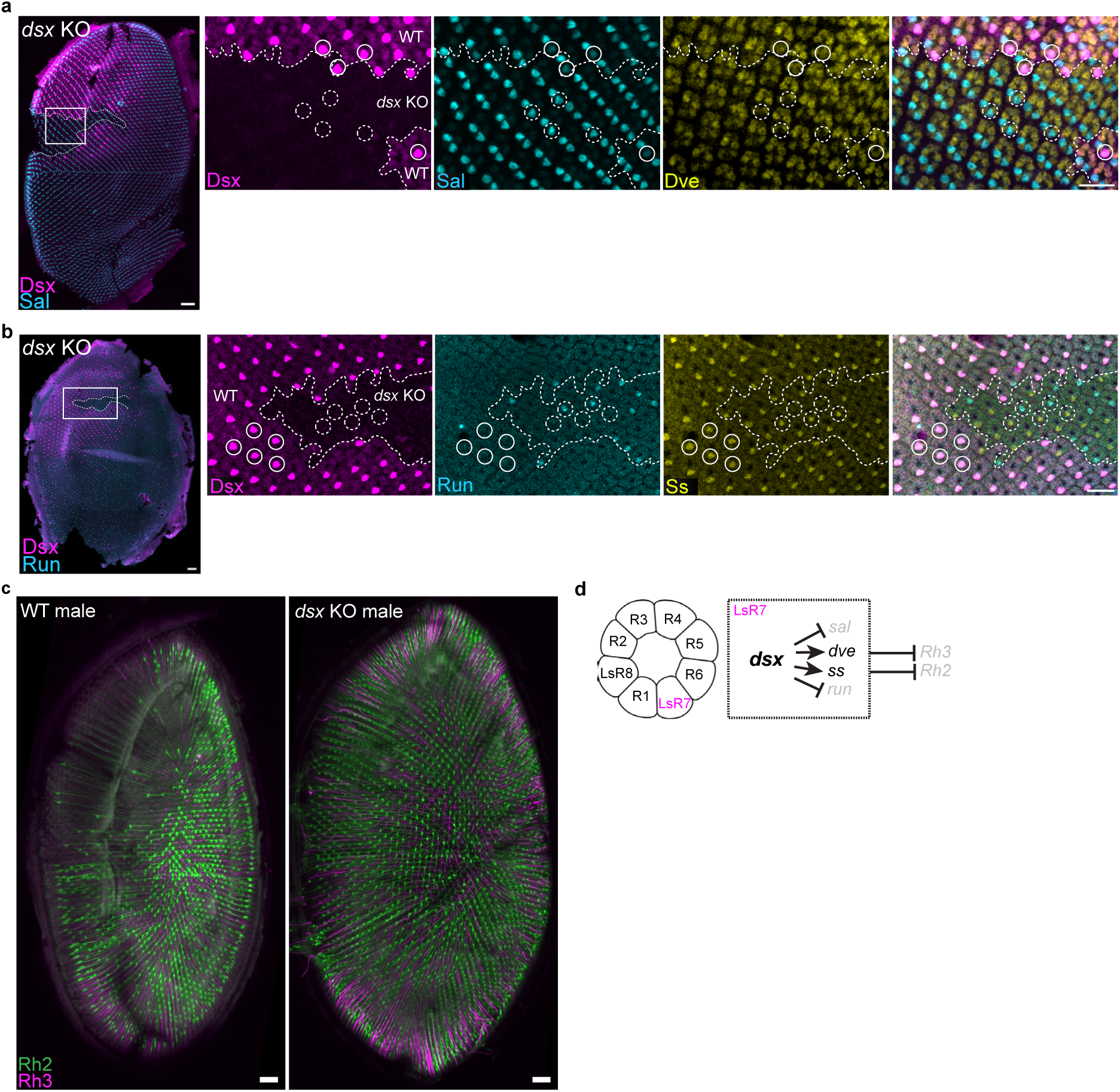
Dsx CRISPR/Cas9 knock-out leads to a loss of LsR7 fate. **a)** Male P50 displaying a *dsx* mutant clone (dashed line) inside the Love Spot. Compared to normal LsR7 (plain circles), *dsx* mutant LsR7 (dashed circles) have a higher Sal level and a decreased Dve level, similar to R8 and non-Love Spot R7. (scale bar whole retina: 50 µm. Scale bar insets 25µm) **b)** Male P50 displaying a *dsx* mutant clone (dashed line) inside the Love Spot. Compared to normal LsR7 (plain circles), which all express Ss and don’t express Run, *dsx* mutant LsR7 (dashed circles) express either Ss or Run, like non-Love Spot R7. (scale bar whole retina: 50 µm. Scale bar insets 25µm) **c)** R7 Rhodopsins expression in WT male adult retina and a *dsx* mutant male adult retina. Loss of Dsx causes Love Spot R7 to express a mix of Rh2 and Rh3 similar to the rest of the retina. (scale bars 50µm) **d)** During pupal development, *dsx* is required in the LsR7 to activate *ss*, upregulate *dve*, repress *run* and down-regulate *sal*. *Dsx* is also required to repress *Rhodopsin 2* (*Rh2*) and *Rhodopsin 3* (*Rh3*).

**Figure 4.**
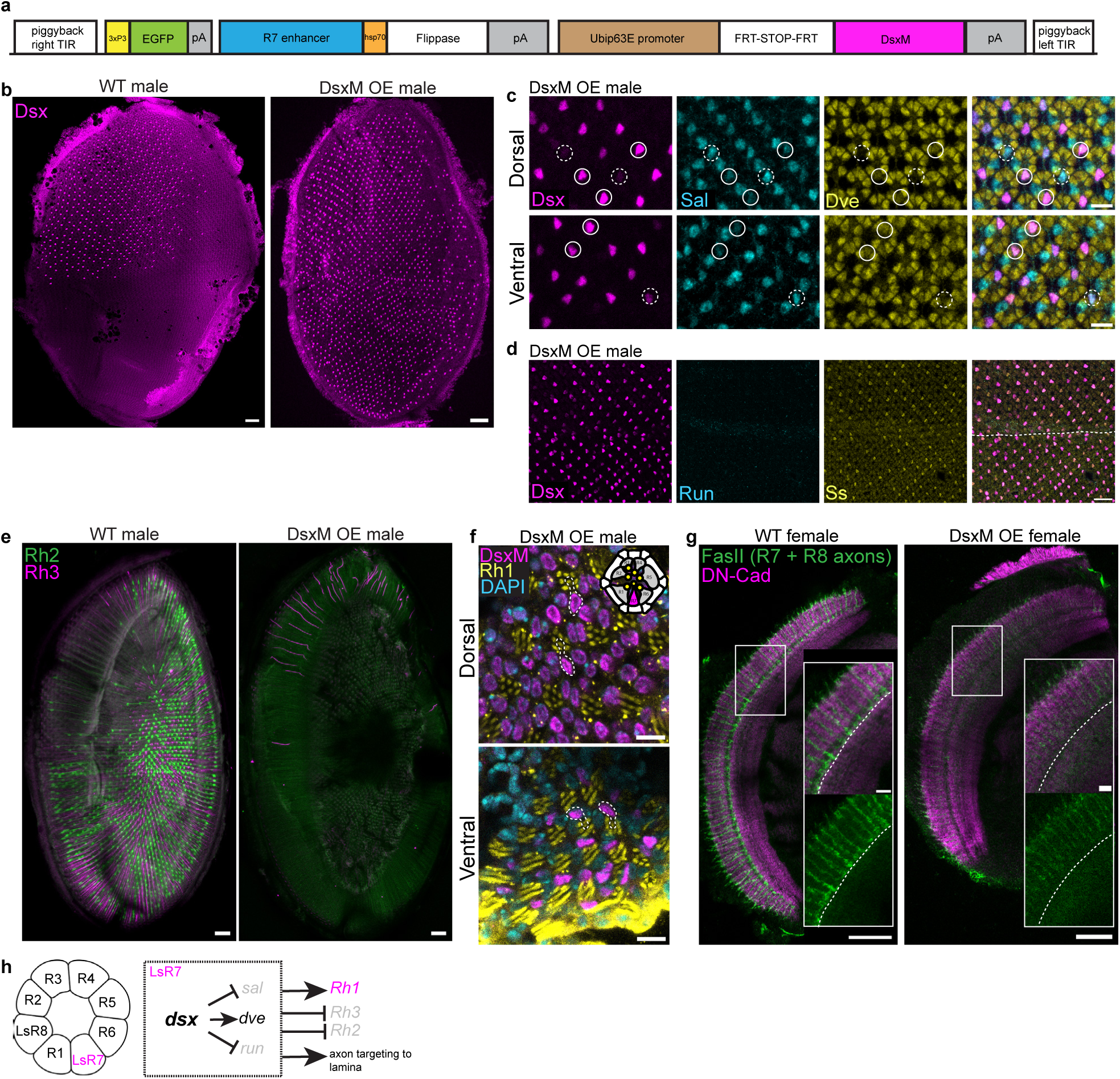
Overexpression of Dsx in the entire retina changes R7 fate to LsR7 fate. **a)** Linear map of the piggyBac construct R7:FLP,Ubi:DsxM used to over-express DsxM in all R7s starting at mid-pupal stages. The insertion marker 3xP3:EGFP is active in the larval brain, which is used for screening. The R7-specific enhancer drives the expression of flippase at mid-pupal stage; flippase then removes the FRT-STOP-FRT cassette which allows constitutive expression of DsxM under the control of the ubiquitin p63E promoter (Ubip63E). These sequences are flanked by piggyBbac right and left terminal inverted repeats (TIR), sequences that are recognized by the piggyBac transposase (encoded on a separate plasmid) to mediate insertion into the housefly genome. Hsp70: *D. melanogaster* heat-shock protein 70 promoter, pA: *D. melanogaster* poly adenylation sequence. **b)** Comparison of Dsx expression in WT and transgenic R7:FLP,Ubi:DsxM flies at P60. The transgenic flies express high levels of DsxM in a majority of R7 across the retina. Scale bar 50µm. **c)** When levels of over-expressed DsxM are high enough (plain circles), DsxM downregulates Sal and upregulates Dve, similar to WT LsR7, in the dorsal and ventral retina. Some R7 express weaker levels of DsxM (dashed circles), which are not sufficient to change Sal or Dve levels. Scale bar 10µm. **d)** DsxM overexpression silences Run in R7s in both the dorsal and ventral eye (dashed line: eye equator) but fails to induce Ss expression in R7. Scale bar 25 µm. **e)** R7 Rhodopsins expression in a WT male adult retina and in DsxM OE male adult retinas. DsxM OE male adult retinas only have residual expression of Rh3 (dorsally, in the dorsal rim area, where both R7 and R8 express Rh3). **f)** Rh1 expression in adult DsxM OE male retinas. DsxM OE adult R7 still over-express DsxM and now express Rh1 (instead of Rh2 or Rh3) like outer photoreceptors. The position of R7 in each ommatidium can be deduced from the geometry of the Rh1-positive rhabdoms (yellow circles, see top-right schematic) and the R7 cell contours are outlined (dashed lines). Scale bar 10 µm. **g)** R7 and R8 axon projections in the medulla of WT and DsxM over-expressing (DsxM OE) adult females (visualized with FasII): in WT, R7 axons terminate in the serpentine layer (black line in the middle of the DN-Cadherin stain) and R8 in a more superficial layer. In a DsxM OE female adult retina, all R7 terminal are missing and only R8 terminals remain visible. Scale bar 50µm, inset: 10µm. **h)** Dsx is able to change R7 fate to LsR7 fate in both sexes and across the retina by repressing *sal*, *run*, *Rh2* and *Rh3*, activating *dve* and *Rh1* and retargeting LsR7 axons to the lamina.

**Figure 5.**
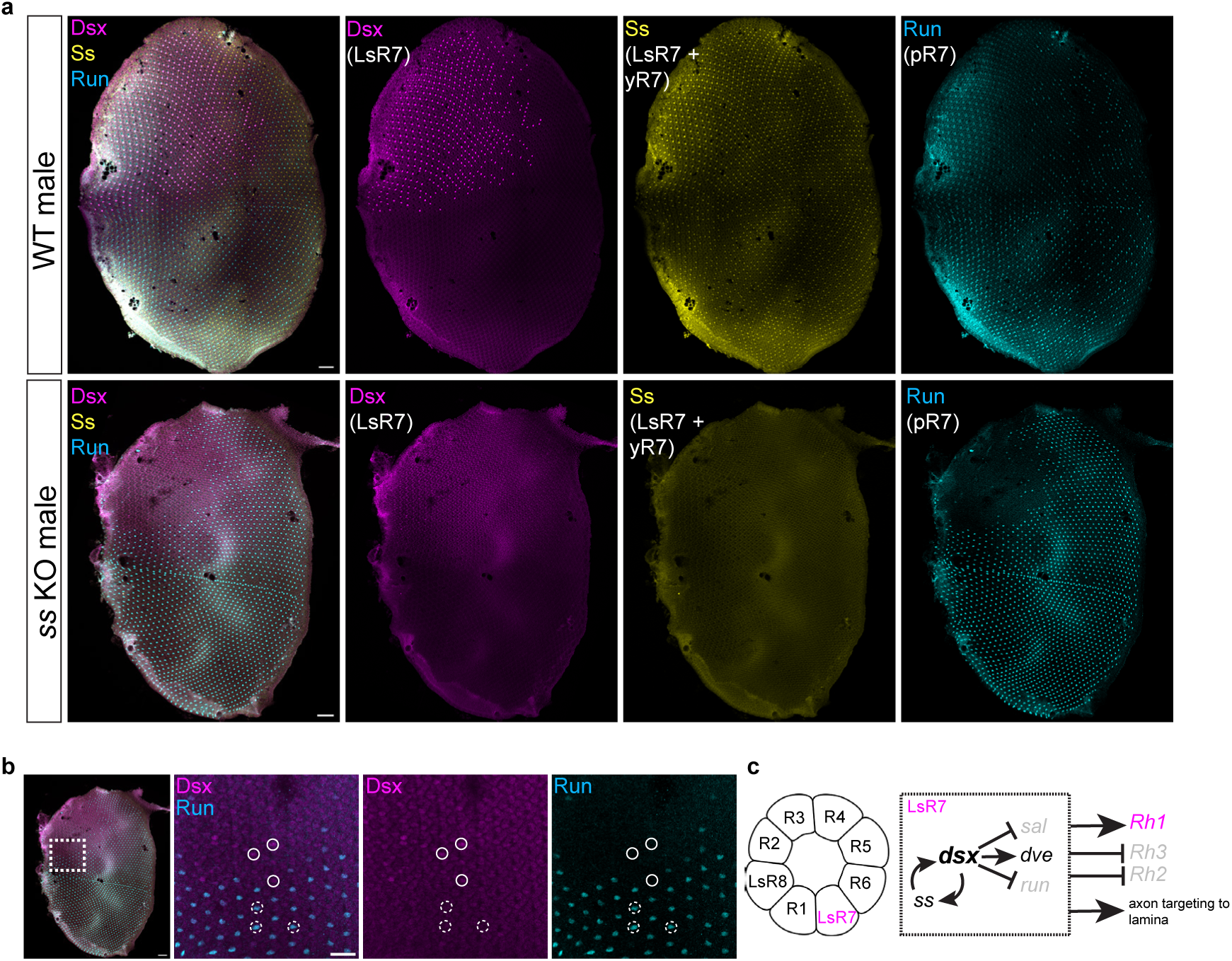
Ss knock out in male pupal retinas. **a)** Compared to WT P50 male retinas (top line), *ss* knock-out male P50 retinas lose the very high levels of Dsx in LsR7. Almost all R7 now express Run (a pR7 marker) except a region at the core of the Love Spot where some Dsx expression persists. Scale bars 50µm. **b)** In *ss* knock-out male P50 retinas, the residual levels of Dsx in some R7s (plain circles) is high enough to suppress Run expression compared to nearby R7s with lower Dsx levels (dashed circles). Scale bar 25µm. **c)** *dsx* interacts with *ss* in a positive feedback loop that strongly activates *dsx* in LsR7. High Dsx levels then repress *sal*, *run*, Rh2 and Rh3, activate dve and Rh1 and retarget LsR7 axons to the lamina.

**Figure 6.**
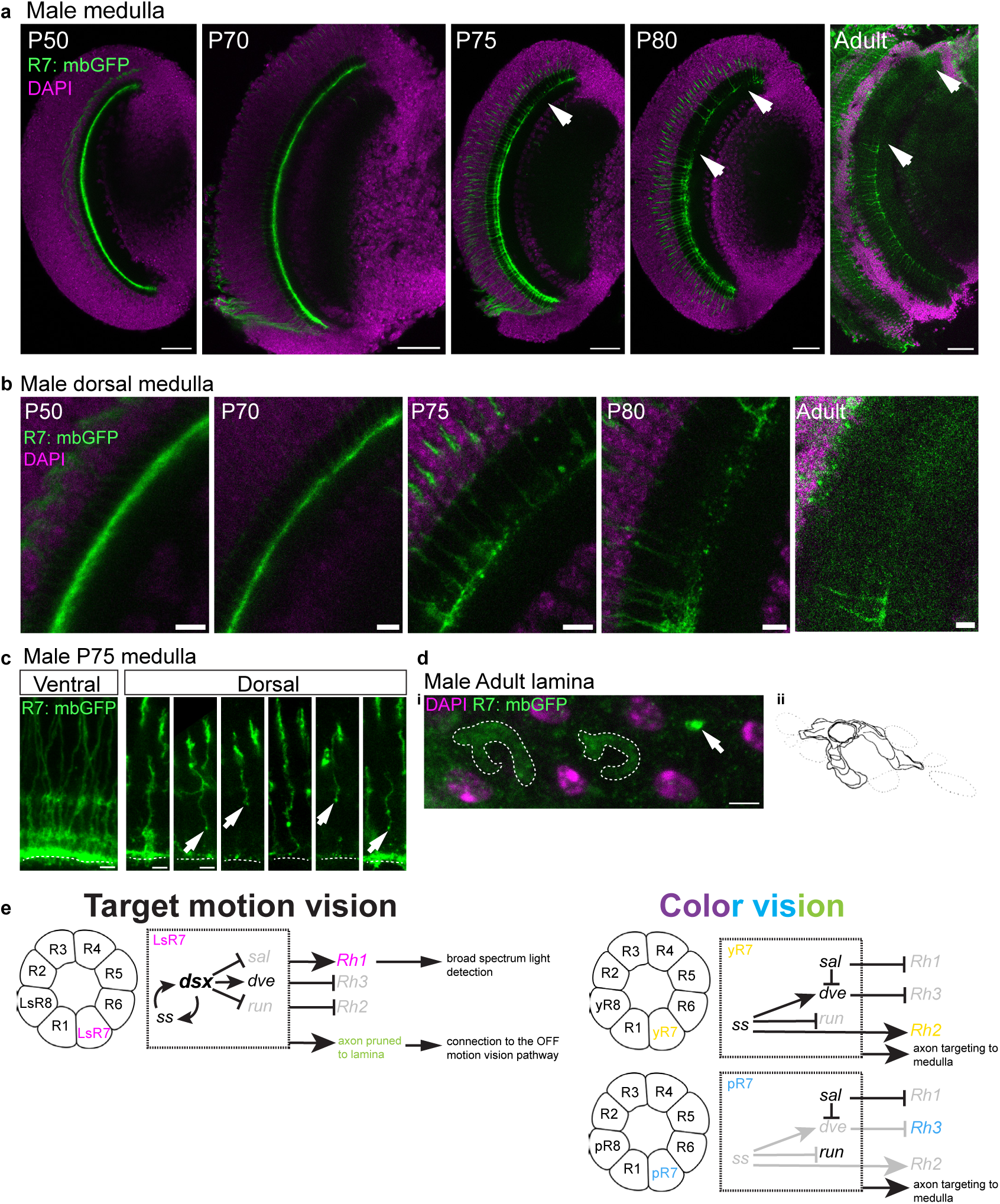
Time course of R7 axon targeting in the male optic lobe. **a)** R7 axon terminal in a R7:mbGFP medulla at different time point of pupal development and in adults. At P50 and P70, all R7 terminals are present in the medulla. At P75, some of the dorsal R7 terminals (ie LsR7 terminals) start missing (arrow), which is even more pronounced at P80 (area between the two arrows). In adults, all dorsal R7 terminals (LsR7 terminals) are missing (area between the two arrows). Scale bars 50µm. **b)** Close-up view of the dorsal region of the medulla shown in A). The continuous line formed by R7 terminals at P50 and P70 starts to break down at P75, when membranous blebs start to appear and axons become thinner and disappear. In adult, dorsal R7 terminal are no longer present in the medulla. Scale bar 10µm. **c)** Close-up view of selected R7 axons from the ventral and dorsal P75 male medulla presented in A). In the ventral medulla, R7 axon terminate around the serpentine layer (dashed line), whereas many dorsal R7 (LsR7) axons are shorter, thinner, and display membranous blebs (arrows). **d)** Bottom view of the distal lamina of an adult R7:mbGFP male, showing the LsR7 terminal and their characteristic lobster-claw shape **(i)**, which had been previously described by electron microscopy in 1983^29^ **(ii).** The axon of a non-LsR7, on its way to the medulla, is also visible (arrow). **e)** Summary of the gene regulatory network in LsR7 compared to yR7 and pR7. In LsR7, *dsx* interacts with *ss* in a positive feedback loop to yield very high Dsx levels. Dsx represses *sal*, activates *dve* (potentially via *sal* repression) and represses *run*. *sal* repression could lead to *Rh1* de-repression, while higher Dve levels could repress *Rh3*. The mechanisms of *Rh2* repression despite the presence of Ss remains unknown but Dsx could repress *Rh2* downstream of Ss. Dsx also triggers LsR7 axon pruning or retraction from the medulla to the lamina, via unknown effectors. These changes in opsin and axon projection allow LsR7 to contribute to motion detection by the Love Spot during male chases. In yellow R7 (yR7) and pale R7 (pR7) the regulation of opsin expression and axon targeting by *ss*, *sal*, *dve* and *run* is likely similar to that known in *D. melanogaster*, which leads to *Rh2* expression in yR7 and *Rh3* in pR7 along with projection of their axon to the medulla, where they contribute to color vision.

To test the role of *dsx* in the divergence of LsR7 from yR7 identity, we knocked it out using somatic CRISPR/Cas9 and assessed photoreceptor differentiation in G0 mutant retinas (**Fig3**). Where *dsx* knock-out (KO) clones, marked by their absence of the protein, intersect the Love Spot region, R7s produce Sal and Dve at levels similar to normal yellow or pale R7s (**Fig3a**). Loss of Dsx also restored stochastic Ss and Run expression, similar to regions outside the Love Spot (**Fig3b**). Interestingly, even hypomorphic clones where Dsx antibody signal is reduced but not gone in LsR7s led to a conversion to a normal R7 fate, including normal stochastic expression of Ss and Run (**Extended Data Fig4**). This indicated that high levels of Dsx are required to turn R7s into LsR7s. Thus, Dsx expression is required for increased Dve, decreased Sal, expression of Ss, and loss of Run in LsR7s (**Fig3d**). In adults, whole-retina *dsx* KO led to a restoration of color vision Rhodopsins Rh2 and Rh3 in a stochastic pattern (**Fig3c**), showing that Dsx is required in LsR7 to change Rhodopsin expression (**Fig3d**).

### Adding target detecting neurons

These results suggested that Dsx acts as a key regulator in LsR7s. Because a hallmark of master regulators is their ability to change cell fate when ectopically expressed, we set out to over-express DsxM (the male isoform of Dsx) in all R7s across male and female retinas using piggyBac transgenesis^46^.

To drive ectopic DsxM expression in R7s we first sought to identify a suitable gene regulatory region using bulk ATACseq on P50 *M. domestica* retinas. A region of accessible chromatin in the first intron of *ss* (in a homologous position to a previously identified R7/R8 enhancer in *D. melanogaster*^47^) drove DsxM expression in all R7s at P50 (**Extended Data Fig5a-c**). However, the activity of this enhancer decreased rapidly after P50, so we next designed a permanent genetic labeling system similar to the “memory cassettes” used in *D. melanogaster*^48^ (**Fig4a**). This approach drove sustained expression of DsxM in both male and female R7s by P60 (**Fig4b**, **Extended Data Fig5d**). DsxM over-expressing R7s showed upregulation of Dve and downregulation of Sal in both dorsal and ventral regions of the eye (**Fig4c**, white circles), compared to a smaller number of R7s that only expressed low levels or no DsxM (**Fig4c**, dashed circles). Furthermore, all R7s with ectopic DsxM lost Run expression (a marker of pR7 fate) (**Fig4d**). Similar results were observed in DsxM over-expressing females (**Extended Data Fig5b,c,g**) Thus, high levels of DsxM in R7s can upregulate Dve, downregulate Sal, and fully repress Run in both males and females (**Fig4h**).

Interestingly, while all R7s that exhibited high ectopic Dsx expression were converted into LsR7s, ectopic DsxM expression at P50 was not sufficient to activate Ss in the stochastic subset outside the love spot that did not already have it, the pR7 (**Fig4d**). In *Drosophila*, the stochastic decision that controls Ss expression is made very early, before differentiation of retina progenitor cells^49,50^, while in this experiment DsxM expression was added after R7 specification. That the addition of DsxM is unable to activate Ss in pR7s suggests that the early expression of DsxM in the future Love Spot region during L3 (**Fig2e**) is required to bias the stochastic expression of Ss to close to 100%. Despite this lack of Ss expression in a subset of R7s, after ectopic DsxM expression we observed upregulation of Dve and downregulation of Sal in both Ss-positive and Ss-negative R7s, demonstrating that regulation of these two genes by Dsx is independent of Ss (**Extended Data Fig5b,c**).

We next assessed the effect of ectopic DsxM on Rhodopsin expression and R7 axon projection as terminal features of LsR7 fate. Adult retinas with ectopic DsxM expression showed an almost complete loss of Rh2 and Rh3 (other than Rh3 in the DRA)(**Fig4e**, **Extended Data Fig5e**), along with a gain of Rh1 expression in the entire retina of both sexes (**Fig4f**, **Extended Data Fig5f**). In addition, upon DsxM overexpression, R7 axon terminals were missing from the medulla, suggesting they were now located in the superficial lamina as in LsR7s (**Fig4g**). DsxM is thus able to upregulate *dve*, downregulate *sal*, repress *run*, and change Rhodopsin expression and axon projection in all R7s in both males and females, demonstrating that DsxM is not only required but fully sufficient to induce LsR7 fate (**Fig4h**).

Interestingly, Ss was not required for the fate conversion to LsR7, which raised the question of its role in wild type LsR7s. In *D. melanogaster*, Dsx has been shown to cross-regulate with cell type-specific factors. For example, in the sex combs, Dsx is initially expressed in a patch on the foreleg in both males and females. In males, DsxM cross-regulates with Sex combs reduced (Scr)^51^. This male-specific feedback loop serves to reinforce *dsx* expression and in turn causes the production of male-specific sex combs^51^. We therefore hypothesized that *dsx* might interact with *ss* in a positive feedback loop in LsR7s.

We used CRISPR/Cas9 to knock out *ss* and examined full *ss* KO male pupal retinas. Strikingly, *ss* KO led to a dramatic loss of Dsx expression in LsR7s (**Fig5a**), showing that Ss expression is required to achieve high levels of Dsx. Interestingly, some residual Dsx expression was still present in R7s of the more dorsal and anterior parts of the Love Spot region and able to repress Run (**Fig5b**). In females, *ss* KO converted all R7s to pR7, Run-expressing fate, as expected (**Extended Data Fig6**). Together with our *dsx* KO results, this shows that Dsx and Ss interact in a positive feedback loop from early stages to produce the very high Dsx levels observed in LsR7 at P50, and that high Dsx levels are required to downregulate Sal, upregulate Dve, change Rhodopsin expression, and alter axon targeting (**Fig5c**).

### Late re-targeting of LsR7 axons

The fact that high DsxM levels in LsR7s are reached late in development, between P30 and P50, raised the question of how Dsx regulates LsR7 axon targeting. *D. melanogaster* R7 terminals reach the medulla shortly after specification in L3 and well before P20^52^, a timepoint at which *M. domestica* LsR7 Dsx levels are still low (**Fig2e**). We speculated that even low early Dsx levels could be sufficient to prevent LsR7 axon from reaching the medulla, causing them to stop in the distal lamina. Alternatively, LsR7 axon could first reach the medulla, like other R7 axons, and then later retract or prune their axon to terminate in the distal lamina.

To visualize LsR7 axons we used the previously identified R7-specific enhancer (**Fig4**, **Extended Data Fig5**) to drive membrane-GFP expression in all R7 starting at mid-pupal stages and followed LsR7 axons during pupal development (**Fig6**). Strikingly, in males, no R7 axons were missing in the medulla at P50 (**Fig6a**), a stage at which Dsx expression in the LsR7 is already very high (**Fig2e**). All axons were still present at P70, and dorsal axons, corresponding to LsR7 axons, only started to go missing at P75, continued to disappear through P80 and were completely gone in adults (**Fig6a,b**). In contrast, in females, all R7 terminals were present in the medulla from P50 to adult stages (**Extended Data Fig7**).

A close examination of the medulla portion of LsR7 axons at P75 revealed that they were thinner and shorter than ventral R7 axons, and displayed membrane blebs reminiscent of observations in axon pruning of *D. melanogaster* mushroom body gamma neurons during metamorphosis^53^ (**Fig6c**, arrows). The axons of LsR7 were later observed in the male adult superficial lamina, with “lobster claw”-shaped feet, similar to the morphology previously described by electron microscopy^29^ (**Fig6d**). Together, these results show that LsR7 axons first reach the medulla and are later pruned or retract back to the superficial lamina.

## Discussion

Drosophila R7 photoreceptor specification is one of the most heavily studied paradigms of cell fate specification. Here we show how this process has been evolutionarily modified to give rise to a new cell type: the LsR7. We characterized this feature using modern tools, examined gene expression during development, and used loss- and gain-of-function approaches to uncover the genetic changes that lead to the specification of this novel neuron. Our results show that Dsx controls the development of a new male-specific photoreceptor type in the housefly Love Spot. We showed that Dsx interacts with Ss in a positive feedback loop and regulates Sal and Dve, along with Rhodopsin expression and axon targeting (**Fig6e**). In *D. melanogaster* retinas, Dve represses Rh3 in outer photoreceptors^44^ while in R7s Sal represses Dve, activates Rh3^44^ and represses Rh1^39^. In yR7s, Ss activates Dve to repress Rh3 while also activating Rh4^54^. This regulatory network is likely conserved in houseflies (**Fig6e**, right). Our data suggest that, in housefly LsR7, Dsx represses Sal, removing Rh1 and Dve repression (**Fig6e**, left). Increased Dve levels in turn repress Rh3 (**Fig6e**, left), while activation of Rh2 by ss is prevented, possibly via the reduction in Sal^43,44^. Further studies could test these regulatory links and assess the direct or indirect nature of Sal and Dve regulation. However, Sal loss in the *D. melanogaster* retina does not disrupt inner photoreceptor axon projection to the medulla^43^: it is thus likely that the effect of dsx on LsR7 axon projection involves other dsx target genes. Potential targets include the ubiquitin-proteasome system, which is involved in *D. melanogaster* mushroom body gamma neuron axon pruning during metamorphosis^53^. That LsR7s first send their axons to the medulla, like other R7s, and later prune them back to the lamina under Dsx control suggests a developmental constraint on the early targeting of R7 axons: levels of Dsx high enough to trigger axon retargeting are reached only at mid-pupation, due to the kinetics of the Ss-Dsx feedback loop, by which point the axons have already reached the medulla.

Although *dsx* knock-out showed that Dsx activates Ss in LsR7, our Dsx R7 over-expression did not lead to Ss activation in all R7. A potential explanation is that Ss activation by Dsx occurs earlier, when the R7 enhancer we use to drive Dsx over-expression is not yet active. Indeed, experiments in *D. melanogaster* show that the stochastic decision of whether to express Ss is made even before retina progenitor cells are recruited as R7 photoreceptors^50^. The early expression of Dsx in retina progenitor cells ahead of the morphogenetic furrow (**Fig2e**) could increase their probability of turning Ss on. Testing this hypothesis would require the identification of a housefly enhancer able to drive DsxM expression in retina progenitor cells anterior to the morphogenetic furrow.

Our *ss* knock out experiment showed that Ss is required for Dsx upregulation in LsR7 at pupal stages. However, the initial cue that activates *dsx* in the presumptive Love Spot region in the larval eye disk remains unknown. Potential candidates are the transcription factors of the Iro-C family (Araucan, Caupolican and Mirror), which are expressed in the dorsal larval eye disc in *D. melanogaster* and are involved in dorsal eye specification^55^.

Other potential Dsx targets in LsR7 are the voltage-gated K+ channel subunits gene *hyperkinetic* (*hk*) and the calcium-activated K+ channel subunit gene *slowpoke* (*slo*), whose LsR7 levels are lower than other R7s and equivalent to those in outer photoreceptors, which could explain why the LsR7 electrical response is faster than other R7s and similar to R1-6^28^. scRNAseq at later pupal stages (after P50) or in adults could reveal additional changes in electrophysiology-related genes in LsR7 that would contribute to the high response speed of Love Spot photoreceptors^25^. Future work could also help address other differences that exist in the Love Spot, such as the change in outer photoreceptor physiology (faster response^25^), as well as the increased lens size and change in R1-6 rhabdom size^21^ that contribute to the higher spatial resolution of the Love Spot^24,25^. These changes likely involve Dsx, which is present at low levels in outer photoreceptors in the Love Spot (see for example Fig4b, WT male retina).

The evolution of the retina Love Spot in housefly also raises the question of the co-evolution of downstream circuits in the optic lobe and central brain. Previous studies in various species have shown that changes in sensory organs can be matched by downstream circuits via neural circuit plasticity. In the case of the Love Spot, based on previous studies in *D. melanogaster*^56^, we anticipate that LsR7 partner Dm8 neurons in the medulla would undergo apoptosis. Other changes in the LsR7 downstream circuit have been described: LsR7 synapse on L2 and L3 OFF-motion detection neurons in the lamina, and the relative contribution of cell-autonomous changes versus cell-plasticity in these new connections remain to be determined. The different morphology of L2 and L3 neurons downstream of LsR7^29^ suggest a cell-autonomous process, maybe through the other regulator of sexual dimorphism in insects, Fruitless. Additional changes of the circuit downstream of LsR7 remain to be explored, in particular the development of male-specific dendritic arborization of lobula ColA and ColB neurons (Col: Columnar), and the development of male-specific ColC, ColD, MLG1, and MLG3 (MLG: Male Lobula Giant) lobula neurons, which receive input from the Lovespot^57^ and could contribute to small target motion detection^38,58^. Other potential candidate neurons downstream of LsR7 include LC11, involved in small dark object detection in *D. melanogaster*^59^, or LC10, involved in female visual tracking in *D. melanogaster* males^60^, both visual projection neurons (VPN) that link the optic lobe to the central brain.

Love Spot-like regions are widespread among the Diptera, with visually striking Love Spots present in at least a dozen families^23,34,61–64^, and can also be found in other insects such as mayflies^65^. Although a systematic characterization of the Love Spot’s phylogenetic distribution remains to be done, it is possible that they evolved many times within the Diptera: studying their development in other species will help us better understand how different species can convergently evolve sexually dimorphic sensory systems.

## Methods

### Housefly husbandry

Wild type *Musca domestica* were obtained from the laboratory of Jeffery Scott at Cornell University. Flies were maintained in culture at 27C in an insectary room in custom-made wire frame and mesh fabric cages (28x28x38cm). Inside the cages, flies were provided with water and a dry mix of milk powder and white sugar. Eggs were collected once a week by putting a small amount of larval food in a paper cup in a cage for a few hours; the eggs and food were then transferred to a 500mL plastic tub filled with larval food and covered with a piece of tissue, and the tub was shaken regularly over the next three days. On day four, larvae were split into two tubs; some were discarded to maintain a medium larval density and fresh food was added along with additional wood chips. Larvae started pupating on day five. Adults eclosed on day eleven and were transferred to new cages.

Larval food was prepared as follows: 250g calf manna, 60g wood chips, 15g dry active yeast, and 900mL DI water were mixed in a plastic container and left at insectary temperature for 15min, after which 600g of wheat bran were added and mixed in. The larval food was kept at 4C for up to a week.

### Single cell RNAseq and single nucleus RNAseq

For scRNAseq, two sets of five male pupal retinas and two sets of five female pupal reinas were dissected in cold PBS and placed in four separate tubes in cold PBS on ice. Pupae were sexed during dissection by examining the shape of developing genitals before removing pupal retinas from each animal. Single cell RNAseq was performed using 10XGenomics single cell 3’ v2 chemistry and sequenced on a Novaseq X. For snRNAseq, 24 retinas from 12 males and 24 retinas from 12 females were dissected in cold PBS, placed in a 1.5mL Eppendorf tube; excess PBS was then removed and the tube snap-frozen in liquid nitrogen before being stored at –80C. The samples were then brought to the UCSD Center for Epigenomics on dry ice. Nuclei were isolated using standard 10xGenomics protocol and processed on a ChromiumX, using the single cell multiome ATAC + gene expression v1. Libraries were then sequenced at the UCSD Institute for Genomic Medicine on a NovaSeq X Plus 25 B. Reads were aligned to the *Musca domestica* genome (GCF_030504385.1) using Cellranger 7.2 on the SDSC supercomputer Expanse. Non-model species like *M. domestica* often have poor genome annotation which can lead to a loss of signal if reads fall out of the existing annotation. In order to limit this loss, we modified the existing annotation file using our previous mapped reads and GeneExt^66^ and re-mapped the reads using the new annotation. The mapped reads were then processed in R v4.5.2 with Seurat v5.4.0. using a standard pipeline (supplementary R script and rds files). To facilitate downstream data exploration, we used a file containing gene orthology relationships between *D. melanogaster* and *M. domestica* (gene_orthologs.gz at https://ftp.ncbi.nlm.nih.gov/gene/DATA/) to edit *M. domestica* gene names to the following format: “Mdom gene ID####Mdom Symbol-Mdom gene name-Dmel ortholog Symbol####Dmel ortholog gene ID”. For example doublesex, previously annotated as “LOC101895413” now appeared as “101895413####LOC101895413-protein*doublesex-dsx####40940” (supplementary python scripts and R script).

### Bulk ATACseq

Ten male pupal retinas were dissected in cold PBS and placed on ice. Nuclei were then extracted using a dounce and processed at the UCSD Epigenomics Core for library preparation and sequencing. Reads were mapped to the Housefly genome with bowtie2 and filtered using scripts provided by R.D. Reed and J. Lewis (Cornell University). Aligned reads were visualized using IGV v2.11.2.

### Micro-injection

Our micro-injection protocol is derived from the protocol described in^46^, with a few modifications, including the use of olive oil, as suggested by the Gompel lab Drosophila germline transformation protocol (http://gompel.org/wp-content/uploads/2021/04/Drosophila-transformation-with-chorion.pdf).

Wild-type housefly eggs were collected by leaving a cup of fresh larval food in a cage for 40min, after which the eggs were transferred and washed in a small cup of water. The floating eggs were then transferred to a small nitex mesh basket with a paintbrush and egg clusters were broken apart with water. The basket was then submerged in a 25% bleach solution for 1min to remove the chorion and then washed with water three times. Eggs were then brought to a 18C injection room, where they were aligned at the edge of a piece of 5% agar gel stained blue with Coomassie blue for improved contrast. Eggs were then affixed to a 1.5 glass coverslip on which edge liquid glue (made by dissolving double-sided tape glue in heptane) had been previously applied and dried. The coverslip was stuck to a glass slide with a drop of water and placed in a small dehydration box containing drierite for 3min. The eggs were then covered with olive oil and injected on an inverted Zeiss Axiophot with aluminosilicate needles (Sutter instrument AF100-64-10) pulled on a Sutter puller P-1000. Tips were broken into sharp points on the edge of the coverslip. After injection, the eggs were left in oil to recover for 15min. The oil was then removed with Chemwipes and the eggs dipped in DI-water for rehydration for 30min at RT. Finally, the coverslips were placed on a 1% PBS/agar gel in a squared Petri dish that was wrapped with parafilm and placed in a humid chamber in our Insectary overnight. The following morning, hatched larvae were picked up with a brush and transferred to a *Drosophila* vial whose walls were previously half-covered with humid chemwipes and half-covered with yeast-paste (made by mixing 8g of dry active yeast with 15mL of Calf manna juice obtained by filtering 50g of calf manna plus 100mL of water through cheesecloth). Yeast paste and water was added once a day until the larvae reach pupation stage, when we added a dry chemwipe at the top of the vial.

### PiggyBac transgenesis

Housefly eggs were injected on their posterior end (round, non-pointy end, future germline) with a mix of donor piggyBac plasmid at 1000mg/mL, helper plasmid coding for a hyperactive version of the piggyBac transposase^67^ at 400mg/mL and 2% cell culture-grade phenol red, filtered on a MilliporeSigma Millex Syringe Filter (Durapore (PVDF), 0.22µm). The following morning, larvae were transferred to a vial that was heat-shocked in a water-bath at 37C for one hour. The larvae were then raised normally to adulthood, when they were transferred to a cage with wild type virgin females. Eggs were collected five days later and newly born larvae were screened for 3xP3:tomato or 3xP3:GFP insertion markers (active in the larval brain^46^), after being immobilized on ice for 10min.

### CRISPR/Cas9 knock-out

gRNAs were designed using CRISPOR (Dsx gRNA1: aatgccgcctgacagccgatcgg, Dsx gRNA2: gcgggtcatggccctgcagacgg, Ss gRNA1: ggacatatctgcgggcaggaagg, Ss gRNA2: aacccgatgtattatccagtagg) Housefly eggs were injected on their anterior end (pointy end, future head) with a mix of high concentration Cas9 protein (PNA Bio CP02) at 900ng/µL, two gRNAs (Synthego) at 900ng/µL each and 1% cell culture-grade phenol red, filtered on a MilliporeSigma Millex Syringe Filter (Durapore (PVDF), 0.22µm). Larvae were reared to pupal stage or adulthood and phenotypes evaluated by immunohistochemistry.

### Immunohistochemistry

Larval eye discs, pupal and adult retinas and brains were dissected and stained as described in^68^. Primary antibodies: chicken anti-GFP (Sigma-Aldrich 06-896, 1/200), mouse anti D. *melanogaster* FasII (DSHB 1D4, 1/10), rat anti D. *melanogaster* DN-Cad (DSHB DN-Ex #8, 1/20), rabbit anti *M. domestica* Rh1a (this study, 1/100), guinea pig anti *M. domestica* Rh2 (this study, 1/100), rat anti *M. domestica* Rh3 (this study, 1/100), rat anti *M. domestica* Rh5 (this study, 1/100), rabbit anti *D. melanogaster* Rh6^69^ (remade for this study, 1/100), rat anti *M. domestica* Dsx-Common (this study, 1/200), rabbit anti *M. domestica* Run (this study, 1/400), guinea pig anti *M. domestica* Ss (this study, 1/200), guinea pig anti *Papilio xuthus* Dve^70^ (1/200), rabbit anti *D. melanogaster* Sal-Common^71^ (remade for this study, 1/200). Secondary antibodies (Alexafluor, 1/400): donkey anti rabbit 647, donkey anti guinea pig 488, donkey anti rat Cy5, donkey anti rat 488, donkey anti rat 647, donkey anti rabbit 647, donkey anti rabbit 555, donkey anti guinea pig 647, donkey anti guinea pig Cy5.

Samples were imaged on a LEICA SP8 confocal microscope with a 40x oil-immersion objective and images were processed with ImageJ.

Immunogens used for antibody production:

*M. domestica* Rh1a: GNGSVTDKVTPDMAC

*M. domestica* Rh2: CMRDDMAEYFKSPRF

*M. domestica* Rh3: NEKAPESASTASTTC

*M. domestica* Rh5: CLGVRERNAAPSVNS

*D. melanogaster* Rh6: HPKYKQVLREKMPCLACGKDDLTSDSRTQATAEISESQA

*M. domestica* Dsx-Common: RQRVMALQTALRRAQQQDEARILQMHEVPPVVHPPTALLNAHHHHHHPLPHHITQQLHHHPHHPHP HLVDVSAVAAAAAAGVGVGPVPPHHIAAAAIPTIRSPPHSDHSANGGGGGGGGGGGGGGSGSGGG GGGSAGGGSNGGGGGVGPSSSSMNGMASSSSAASSSTAPPHHTPPDHTHHHHHHHHPHPHLVSV PPTAQSVDSSCDSSSPSPSSTSGVAVPVLVPNRKPNPEQQQNGADMSIDLILDYCQKLIEKFGYPWE MMPLMYVILKDAGVDIDEASKRIEE

*M. domestica* Run: MHLPAGPTMVAAPAQALPSSNNNNSNNNRANSASSTGSSTPDNTSNAAKMPSSMTDMFASLHEMLQ EYHGELAQTGSPSILCSALPNHWRSNKSLPGAFKVIALDDVPDGTLVTIKCGNDENYCGELRNATAIMK NQVAKFNDLRFVGRSGRGKSFSLTITIATYPVQIASYTKAIKVTVDGPREPRSKQSYGYPHPGAFNPFM LNPAWLDAAYMTYGYADYFRHQAAAAAVHHPALAKTSPSLPNGQTVVPPPQAAAAGSPNEYRLPSQI TPPPSGVNAIPATSMIPSPPGSTYSMPQFPFNPVAAAAAAAAASAHPDLLHPHQKPAHAHAFHPYNLA ALRGGRPSHLHNTSLGSHSSTEHISPAHISPASSRPSSSSPTHQALHNKLNTSVESNSLHEQSASDAD SDDEQIDVVKSAFVPILRPQVPSLNGSTEDLDKSLDSARSSPVQPLGRQRCDLKAPSAMKPYYHESS SRQSTSPETTLPAATKLKNASSQQKTVWRPY

*M. domestica* Ss: QQPSQSHSSAAVSPHDQHHHPAQHHSAHHHHHHHHHATHGSHHPHPINHHHHHAAGSAASNSAAT LPTTAAATNGSEVWSPASYASQYASQYFSYHPHHHHHHSHHPHSAAHHSLHAAGSSISTAAPPPVLP PSALATQPPAPLPPHISSSTQ

*D. melanogaster* Sal-Common: MNSLDLLQKRAQEVLDSASQGILANSMADDFAFGEKSGEGKGRNEPFFKHRCRYCGKVFGSDSALQI HIRSHTGERPFKCNVCGSRFTTKGNLKVHFQRHAQKFPHVPMNATPINTCETMKLKELMKNKKISDP NQCVVCDRVLSCKSALQMHYRTHTGERPFKCRICGRAFTTKGNLKTHMAVHKIRPPMRNFHQCPVC HKKYSNALVLQQHIRLHTGEPTDLTPEQIQAAEIRDPPPSMMPGHFMHHHHHH

### Molecular biology

Genomic DNA extraction was performed on wild-type pupae as described in ^64^.

PCR was performed using NEB Q5 DNA polymerase and DNA fragments were purified using a PCR cleanup kit or gel extraction kit (Qiagen). Gibson primers were designed with Benchling assembly wizard and Gibson assembly was done using NEB HiFi DNA assembly master mix.

The piggyBac helper plasmid with the *D. melanogaster* codon-optimized hyperactive transposase was a gift from the Wimmer lab (Georg-August-University Goettingen, Germany).

The R7:DsxM plasmid was built by Gibson assembly from p755 (Akbari lab), a commercially synthesized *M. domestica* DsxM (Genescript) and the R7 enhancer (amplified from *M. domestica* genomic DNA with forward primer tctcctgggcctcatatatgtctcc and reverse primer tctctctctgtttggacaaattgttttaaaagg)

The R7:FLP,Ubi:DsxM piggyBac plasmid was built by Gibson assembly from the R7:DsxM plasmid cut with NheI (NEB), *M. domestica* Ubip63E promoter (amplified from *M. domestica* genomic DNA with forward primer ctgctgctttgcattgttccttcac and reverse primer gactaagctagtttgtcagtctaaagaaaggtagaatg), flpD5 coding sequence (amplified from YZ23^50^) and FRT-STOP-FRT (amplified from Addgene plasmid #71809).

The piggyBac 3xP3-tdtomato plasmid was built by replacing GFP in PL755 with tdTomato from PL1017A (Akbari lab, UCSD) via a PCR and Gibson assembly.

The R7:CD4-GFP was built by Gibson assembly from our piggyBac 3xP3-tdtomato cut with FseI and NheI (NEB), the R7 enhancer (amplified from *M. domestica* genomic DNA) and CD4-tdGFP (amplified from Addgene plasmid #31221).

All plasmids used for microinjection were transformed into NEB stable competent cells (made with Zymo kit) and prepared using a Qiagen midiprep kit.

### Focus stacking pictures of housefly heads

Photos were taken on a Nikon z7 camera with a 10x microscope objective (Mitutoyo) on a custom focus-stacking set up. The photos were taken with a 25µm increment and merged using Zerene Stacker software using the Pmax algorithm.

### Generation of retina schematics (Fig1e)

A grid of randomly arranged yellow and blue hexagons was generated with a custom python script using the package hexalattice. The grid was then cropped to resemble a fly eye and the DRA and the Love Spot hexagon colors were edited in Inkscape.

## Accession codes

### Primary accessions

#### Sequence Read Archive

SRPxxxxxx – Data will be deposited upon or before manuscript acceptance

#### Data deposits

Data will be deposited upon or before manuscript acceptance at SRA under accession code SRPxxxxxx.

## Acknowledgements

We thank Claude Desplan and members of his lab, Mathias Wernet, Gregor Belušič, Neset Ozel for helpful feedback and discussion, Bogdan Sieriebriennikov and James Lewis for help with bioinformatic analysis. This work was supported by a Hellman Fellowship (Hellman Fellows Fund, San Francisco, CA), NIH NHGRI grant R01HG013634, and AFOSR grant FA9550-23-1-0530 to M.W.P.

## Contributions

M.P. and A.D. conceived the project; A.D. and M.P. designed experiments and A.D., M.P., N.K., Y.Z. and

E.T. performed experiments. A.D. and N.K. analyzed single cell data. M.P. supervised the project. M.P. and A.D. wrote the paper.

## Ethics declarations

The authors declare no competing financial interests.

---

**Extended Data Figure 1.**
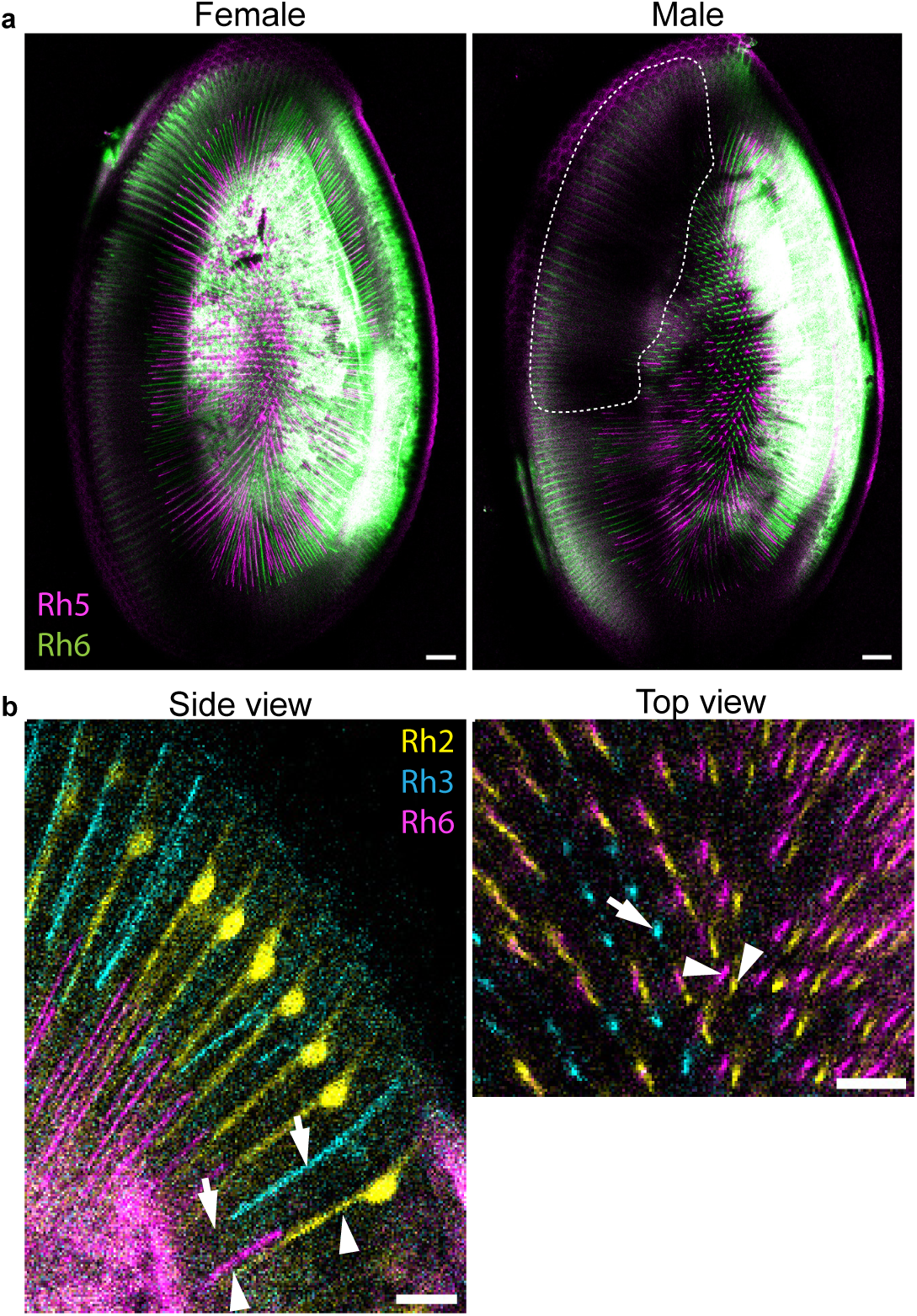
Opsin expression in R8. **a)** Female and male adult housefly retina R8 express either Rhodopsin 5 (Rh5, magenta) or Rhodopsin 6 (Rh6, green). In the Love Spot (Male retina dotted area), R8 express neither Rh5 nor Rh6. Scale bar 50µm. **b)** Rh6-expressing R8 lie under Rh2-expressing yR7s (arrowheads), while Rh5-expressing R8 lie under the remaining Rh3-expressing pR7s (not stained here for antibody host compatibility reasons) (arrows). Scale bars 20µm.

**Extended Data Figure 2.**
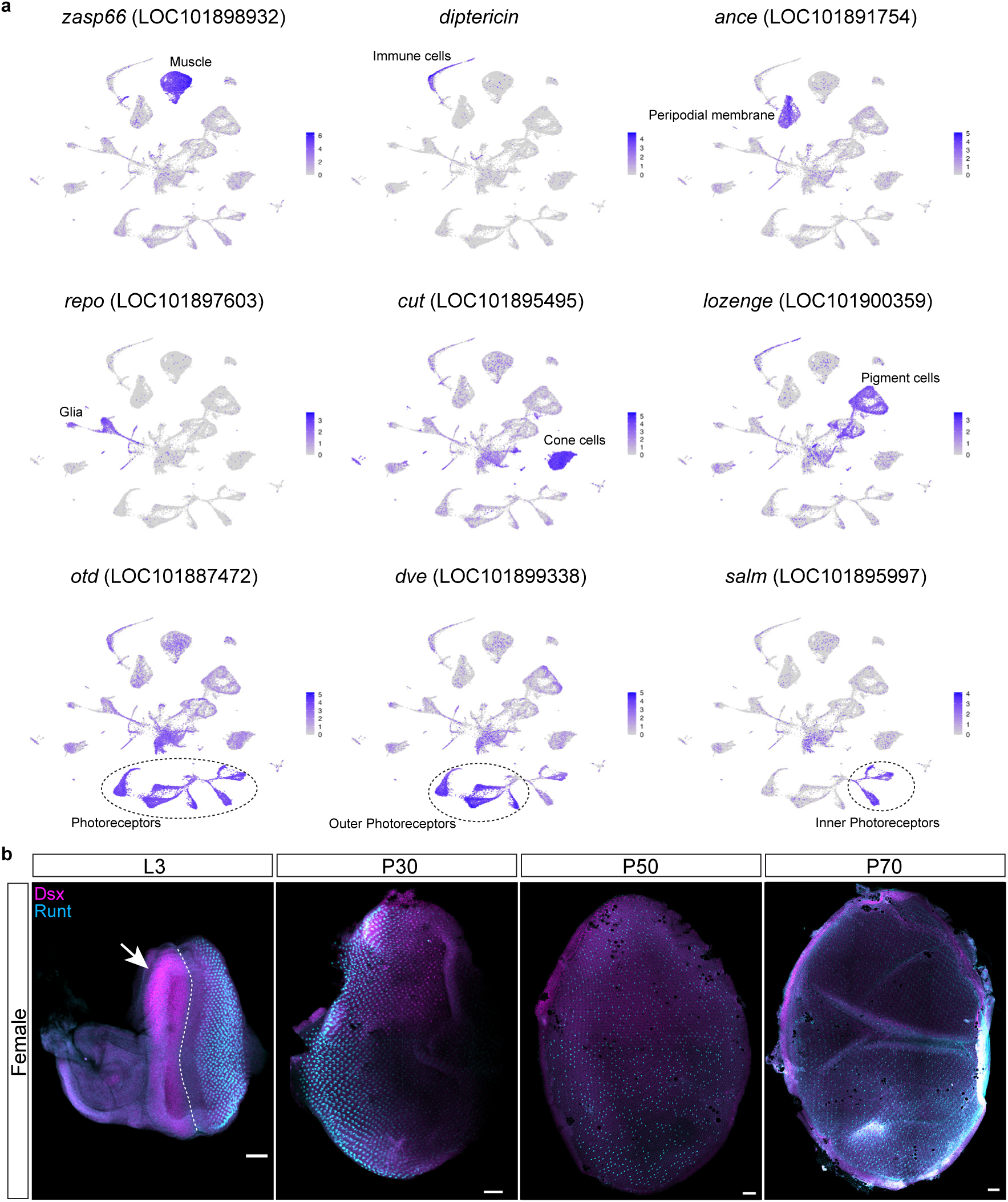
Expression of marker genes used to identify pupal retina cell types in Fig2A and expression of Dsx in the developing female retina. **a)** Gene names are displayed above each UMAP plot along with the locus number (LOCXXXXXXXXX). Many of the observed cell clusters could not be identified precisely (in part because a similar dataset for *D. melanogaster* has not yet been published), but we could still assign most of them to different cell-type families based on the expression of a few marker genes: 4 glial cell clusters expressed *repo* or *wrapper*. A cluster of muscle cells expressing myosin and actin-related genes (such as *zasp66*) could correspond to the retinal muscle recently shown to have a role in vision in *D. melanogaster*^72^. Another cluster was enriched in immune-related genes such as *diptericin* (coding for an antibacterial peptide), whereas a cluster expressing *ance* likely corresponds to peripodial cells surrounding the pupal retina^41^. Cell types making up the ommatidia were also identified based on marker genes known in *D. melanogaster*: cone cells expressed *cut*, pigment cells *lozenge,* and photoreceptors *otd*, inner photoreceptors (R7-8) *salm* and outer photoreceptors (R1-6) *dve*. **b)** At late larval stages (L3), Dsx is also expressed in a dorsal region of the eye disk, before the morphogenetic furrow (dashed line) in females (arrow). By P30, Dsx expression is ramping up in a few dorsal-anterior R7s but remains weak. At P50 and P70, Dsx levels are weak across the retina. Scale bars 50µm.

**Extended Data Figure 3.**
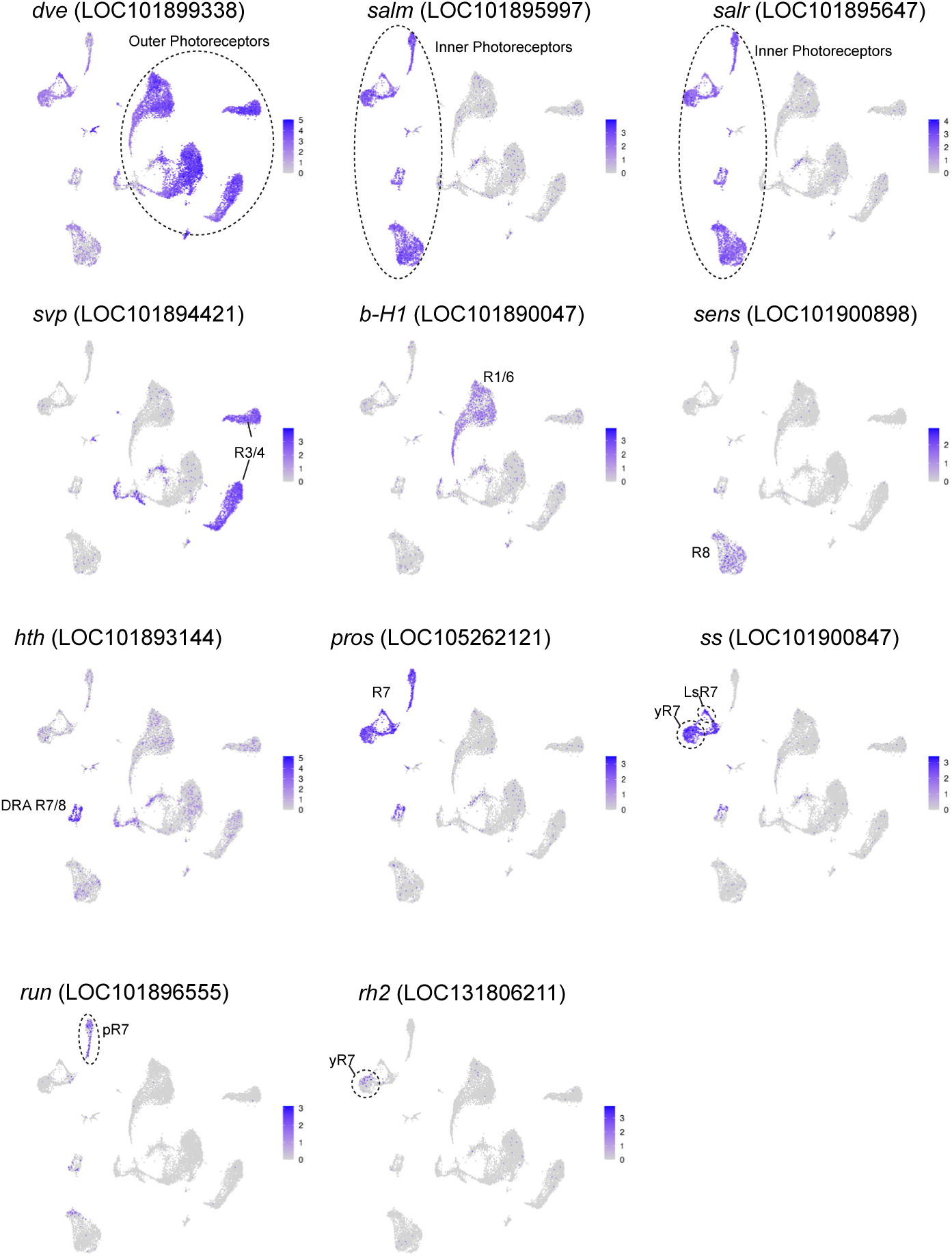
Expression of marker genes used to identify pupal retina photoreceptor types in Fig2B. **a)** Gene names are displayed above each UMAP plot along with the locus number (LOCXXXXXXXXX). Outer Photoreceptors: R1-6, Inner Photoreceptors: R7-8. DRA: Dorsal rim area. yR7: yellow R7, LsR7: Love Spot R7, pR7: pale R7.

**Extended Data Figure 4.**
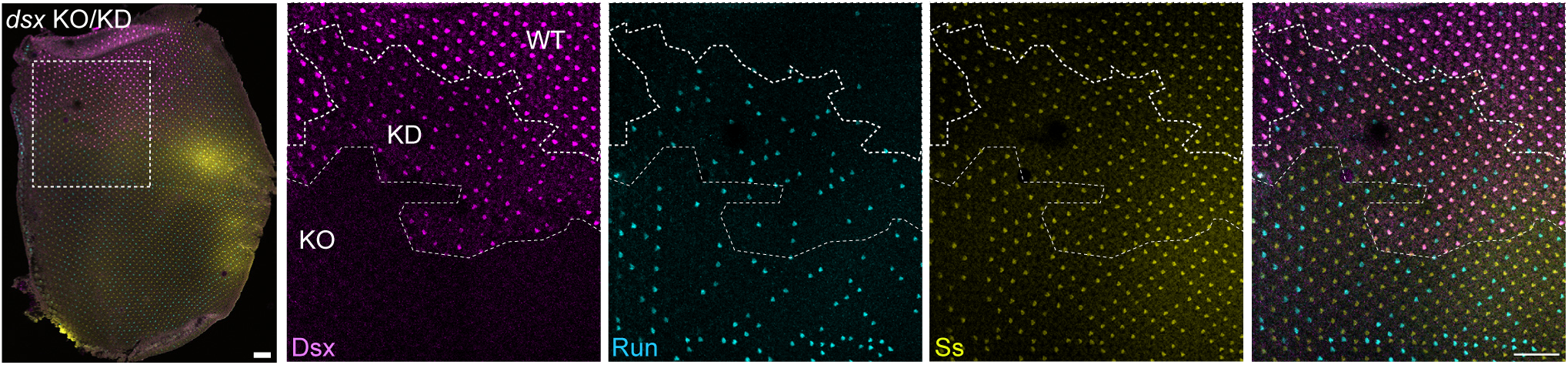
Dsx knock down is sufficient to bring back normal Ss and Runt expression in R7. **a)** A P50 male retina displaying a dsx knock out (KO) clone and adjacent knock down (KD) clone in the Love Spot. In contrast to the WT Love Spot, where all R7 express Ss and not Run, dsx KD R7 and dsx KO R7 express either Ss or Run.

**Extended Data Figure 5.**
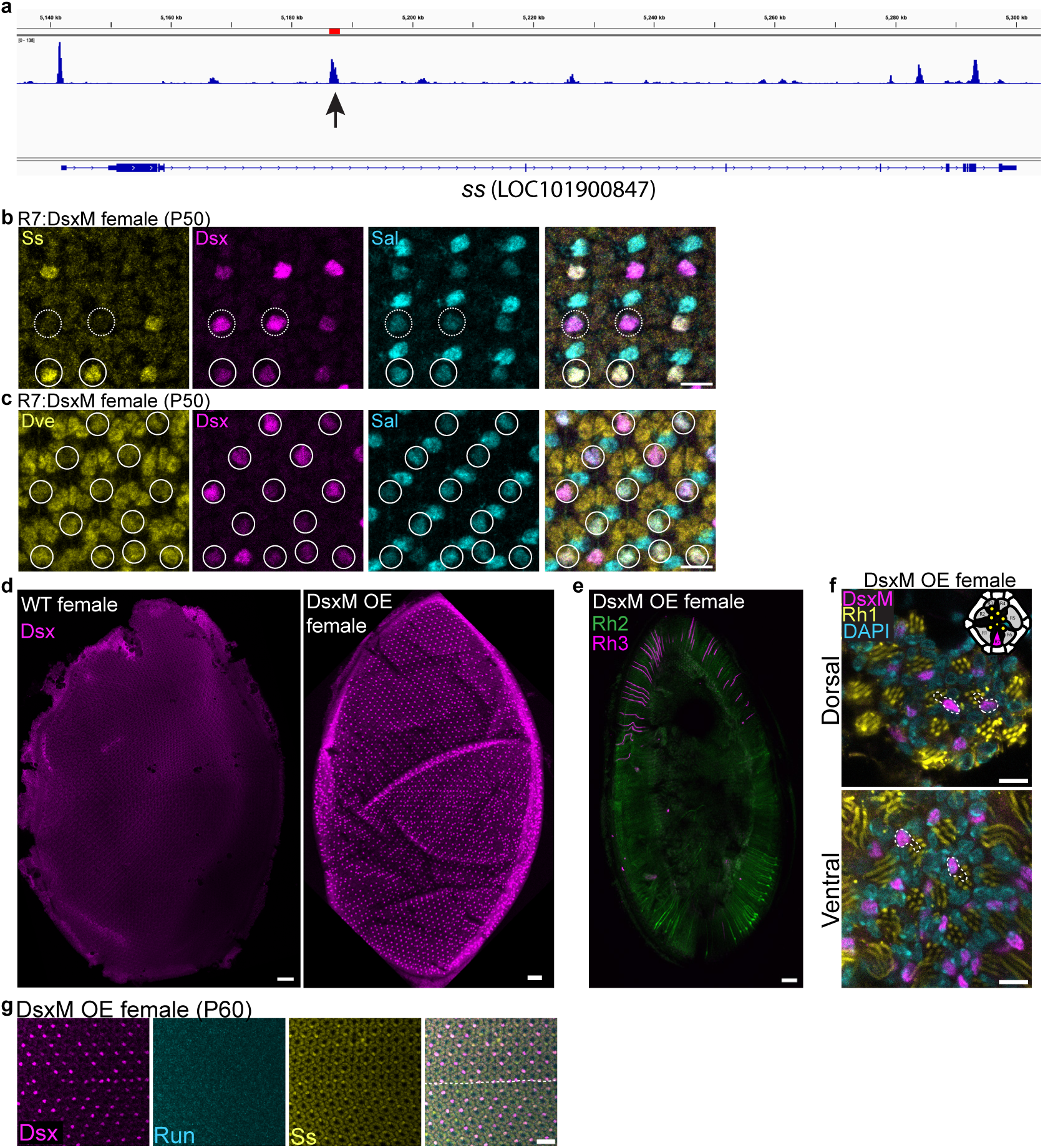
R7 enhancer genomic location and ability to drive DsxM over-expression in all R7. **a)** *Musca domestica* R7 enhancer (red segment) is located in *ss* intron 2, at the location of an ATACseq peak (arrow) (male P50 retinas bulk ATACseq). **b)** Driving DsxM expression in all R7 with the R7 enhancer in female transgenic houseflies leads to a reduction of Sal expression in both yR7 (expressing Ss, plain circles) and pR7 (not expressing Ss, dashed circles), showing this regulation is independent of Ss. **c)** Driving DsxM expression in all R7 with the R7 enhancer in female transgenic houseflies leads to an increase in Dve expression in all R7(plain circles), showing this regulation can happen in both yR7 and pR7 and is thus independent of Ss. **d)** Comparison of Dsx expression in WT and transgenic R7:FLP,Ubi:DsxM female flies at P60. The transgenic flies express high levels of DsxM in a majority of R7 across the retina. Scale bar 50µm. **e)** R7 Rhodopsins expression in a DsxM OE female adult retinas. DsxM OE male adult retinas only have residual expression of Rh2 (ventral retina, green) and Rh3 (dorsally, in the dorsal rim area, where both R7 and R8 express Rh3). **f)** Rh1 expression in adult DsxM OE female retinas. DsxM OE adult R7 still over-express DsxM and now express Rh1 (instead of Rh2 or Rh3) like outer photoreceptors. The position of R7 in each ommatidium can be deduced from the geometry of the Rh1-positive rhabdoms (yellow circles, see top-right schematic) and the R7 cell contours are outlined (dashed lines). Scale bar 10 µm. **g)** DsxM overexpression in female dorsal and ventral P60 retinas R7 silences Run but fails to induce Ss expression (dashed line: eye equator). Scale bar 25 µm.

**Extended Data Figure 6.**
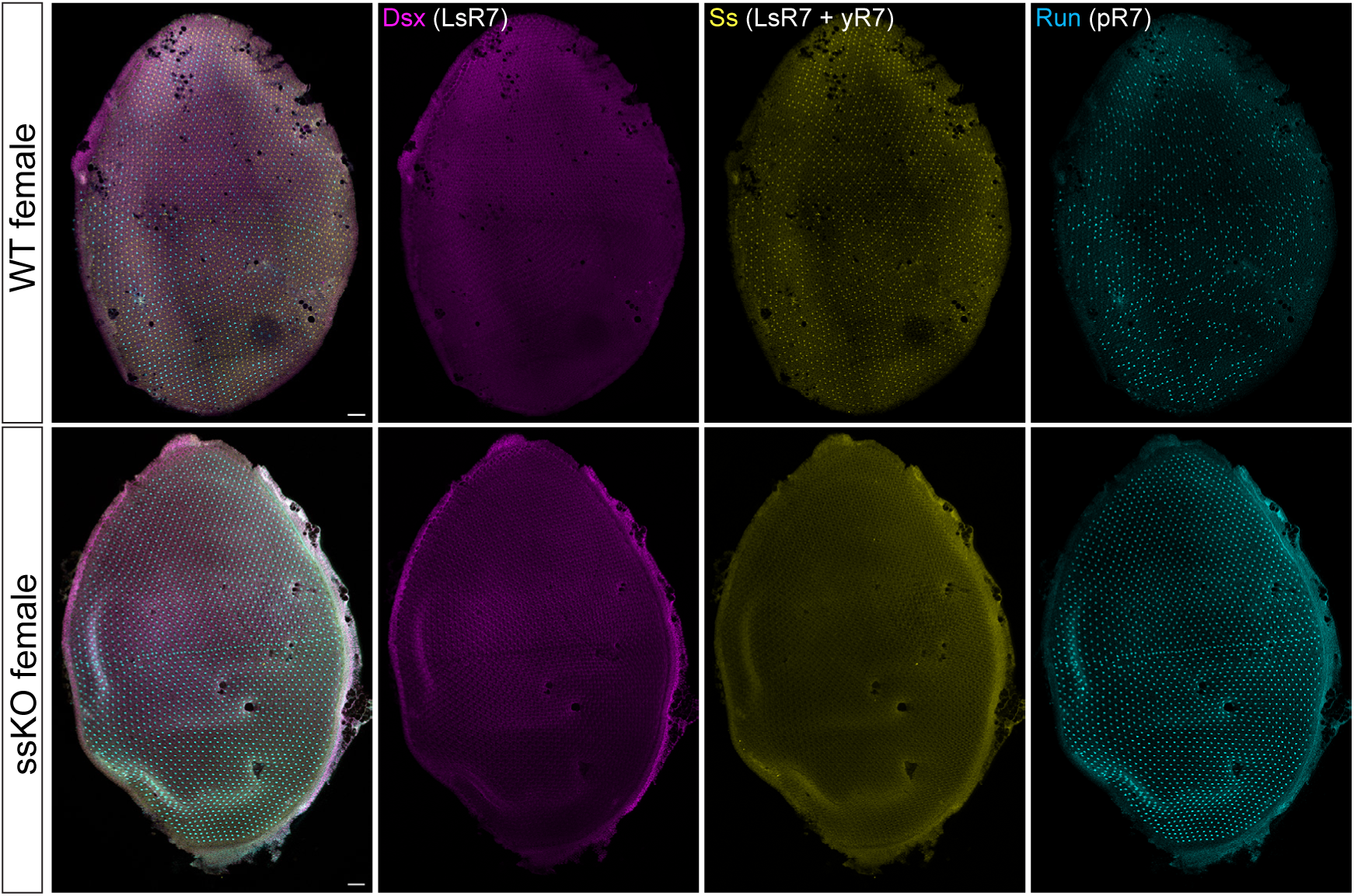
*Ss* knock out in female pupal retinas. WT female P50 retina (top row) showing broad, weak levels of Dsx expression and a stochastic mosaic of Ss and Run-expressing R7. *Ss* knock out leads to expression of Run in all R7s but does not affect Dsx.

**Extended Data Figure 7.**
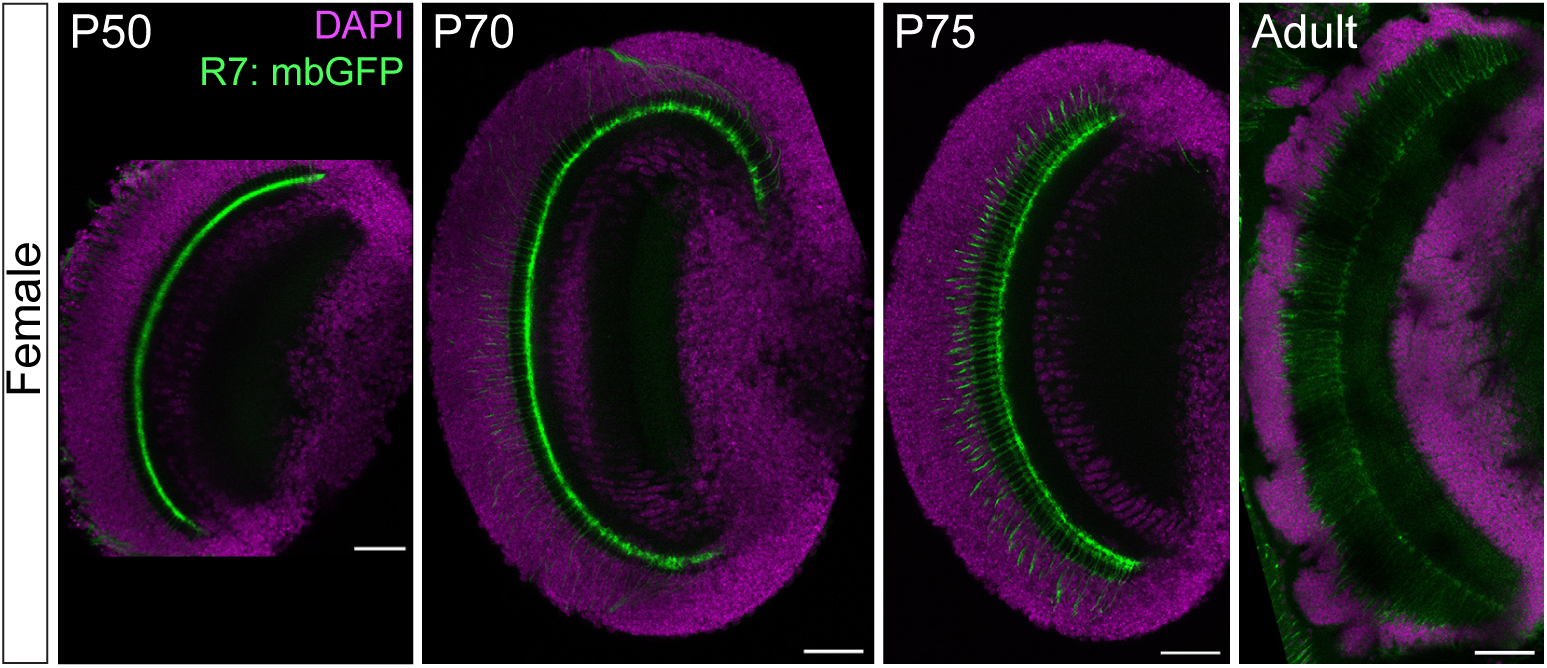
Timecourse of R7 axon targeting in the female optic lobe. The R7 axon terminals in this R7:mbGFP female are all present in the medulla at P50 and remain present across pupal stages and in adults.

